# Reconcile sensory attenuation and enhancement: The temporal dynamics of self-generated sensory feedback

**DOI:** 10.1101/2024.10.07.616960

**Authors:** Yosuke Suzuishi, Acer Yu-Chan Chang, Wen Wen

## Abstract

Self-generated touches are often perceived as weaker than externally generated ones, a phenomenon known as sensory attenuation. However, recent studies have challenged this view by providing evidences that actions could also enhance predicted touch. To investigate this paradox, we examined the temporal dynamics of perceptual processing using steady-state somatosensory evoked potentials (SSSEP) and sliding time window analysis. Results showed that SSSEP in the very early window was smaller in the no-delay and active conditions compared to the delayed and passive condition, respectively, consistent with sensory attenuation. However, this attenuation soon disappeared and reversed in the later windows and onward, indicating an early attenuation followed by a later enhancement. This study resolves the paradox of sensory attenuation and enhancement in self-generated sensory signals by demonstrating that these processes occur at different phases of sensory processing. The findings reveal a sophisticated dynamic balance in sensory processing, facilitating optimal interaction with the environment.

## Introduction

In our daily lives, we receive various types of information, not only from external sources but also from ourselves. For example, we often rub our fingers on a surface and use the information from active touches to infer the texture of the surface. It has been widely reported that self-generated sensory feedback is usually perceived as less intense than externally generated feedback, a phenomenon known as sensory attenuation (Hughes et al., 2013; Wolpert, 1997). For instance, self-generated touches are typically perceived as weaker than those generated externally (Blakemore et al., 1998). Similarly, self-generated sounds trigger smaller amplitudes of event-related potentials (ERP), such as the N1, compared to externally generated sounds (Baess et al., 2011; Gentsch & Schütz-Bosbach, 2011; Kühn et al., 2011; Pinheiro et al., 2019; Timm et al., 2013)

When people perform voluntary actions, the brain is thought to generate predictions of sensory input using efference copies of motor commands, which may ‘cancel’ the anticipated sensations (Wolpert, 1997). The neural basis of this mechanism is known as corollary discharge (Crapse & Sommer, 2008; Subramanian et al., 2019). This mechanism helps the brain distinguish between sensory stimuli generated by our own actions and those originating from the external environment. For instance, the difficulty in tickling oneself is attributed to the brain’s inhibition of self-generated sensations (Blakemore et al., 1998, 2000). The theory of sensory attenuation and corollary discharge aligns with symptoms observed in patients with psychosis, such as schizophrenia. These patients often exhibit a diminished ability to distinguish between self-generated and externally generated sensory input (Frith, 2000; Waters et al., 2012). They also show reduced sensory attenuation (Ford et al., 2007; Heinks-Maldonado et al., 2007; Whitford, 2019). Furthermore, individuals scoring high in schizotypal personality traits report that self-generated and externally-generated tactile stimulation are equally ticklish (Lemaitre et al., 2016).

However, is self-generated sensory feedback always attenuated? This remains controversial. In psychological studies, self-relevant information is reported to usually receive more cognitive resources and is processed more precisely. For example, self-faces and self-names automatically capture attention, i.e., the so-called cocktail party phenomenon (Cherry, 1953; Wolford & Morrison, 1980; Wood & Cowan, 1995). Furthermore, self-generated sensory feedback can be predicted by our brain. It could also lead to a preactivation of the neurons responsible for the sensory processing of the expected input (Roussel et al., 2013). Therefore, the sensory processing of self-generated (and self-relevant) sensory information can potentially be enhanced compared to externally generated (non-self-relevant) sensory input at certain points. Moreover, there are also studies that report the opposite phenomenon of sensory attenuation in specific conditions (Press et al., 2020; Thomas et al., 2022). Particularly, Thomas et al. (2022) reported that when removing the tactile input from the acting finger (i.e., the finger that pushed the button), the perceived intensity of the touch was stronger in active trials than in passive trials. Another later study suggested that this enhancement may be driven by the reference condition used, rather than being a direct result of action (Job & Kilteni, 2023). Yet, another possibility exists: When removing the tactile input from the surface of the acting finger, the timing of the expected sensory input becomes ambiguous, diminishing the sensory attenuation phenomenon and even causing the reversed effect. The temporal predictability of the self-generated sensory input may be critical for the phenomenon of sensory attenuation (Pinheiro et al., 2019) In other words, self-generated sensory attention and enhancement may be distributed at different temporal scales.

To examine the above hypothesis, we combined the steady-state somatosensory evoked potential (SSSEP) and sliding window analysis for a 2-s 71-Hz vibration delivered to one’s finger after a self-paced key press. In Experiment 1, the vibration was either delivered immediately after the key press or after a random delay of 500-1000 ms. In Experiment 2, the vibration was either delivered by the participant’s key press or passively received. In both experiments, participants were asked to judge whether the vibration felt stronger or weaker than the standard stimulation delivered 500 ms after the first stimulation. The SSSEP amplitude in each 1-s time window, which was shifted every 100 ms for the first vibration, was compared between the conditions in each experiment.

## Methods

### Participants

Nineteen participants (6 females, 1 left-handed, mean age = 25.47, range 18 - 38) participated in Experiment 1, and 18 participants (13 females, 1 left-handed, mean age = 20.89, range 18 - 31) participated in Experiment 2. A total of 18 participants and 17 participants completed the main task in Experiment 1 and Experiment 2, respectively, and were included in the data analysis. One participant was excluded from each experiment due to the failure to establish the 50% discrimination threshold for the standard stimulus (see below). The sample size for Experiment 1 was determined using a power calculation conducted with G*Power 3 (Erdfelder et al., 2009; Faul et al., 2007) to ensure a power of 0.80 with an effect size of dz = .72 for the difference in SSSEP between the no-delay and delayed conditions. The sample size for Experiment 2 was based on the effect size of the first time-window in Experiment 1, aiming to ensure a power of 0.95. All participants reported normal motor ability and normal tactile sensation. The study was approved by the local ethics committee (approval code: 23-06). All participants provided written informed consent before participation and received 2000 Yen as reimbursement.

### Stimuli and Apparatus

Tactile vibrations at 71 Hz were applied using an 8-pin Piezostimulator (QuaeroSys) taped to the participants’ left-hand index finger. The Piezostimulator is a mechanical tactile stimulator that drives the pins to different heights between 0 and 1.5 mm with a timing accuracy of 0.5 ms (i.e., the possible stimulation frequency is up to 2000 Hz). The intensity of the tactile stimulus was adjusted by modifying the height of the pins. The stimulation frequency was selected based on pilot experiment results to ensure large and clear SSSEP. Multiple frequencies in the range of 20-100 Hz were tested, and the selected frequency of 71 Hz produced the clearest SSSEP peak.

During the experimental tasks, participants placed their left hand on a table inside a box with their palm facing up. A keypad was placed on the box, ensuring that the key was directly above the tip of the index finger (**Figure 1**). Participants were instructed to press the key with their right index finger to trigger the tactile stimulus during the task. For the intensity judgment, participants responded using two foot pedals (left pedal: the first stimulus felt stronger; right pedal: the second stimulus felt stronger). The experiment was programmed using Python with PsychoPy (Peirce, 2007, 2008) Instructions and a fixation dot were presented in white on a display with a gray background.

**Figure 1.**
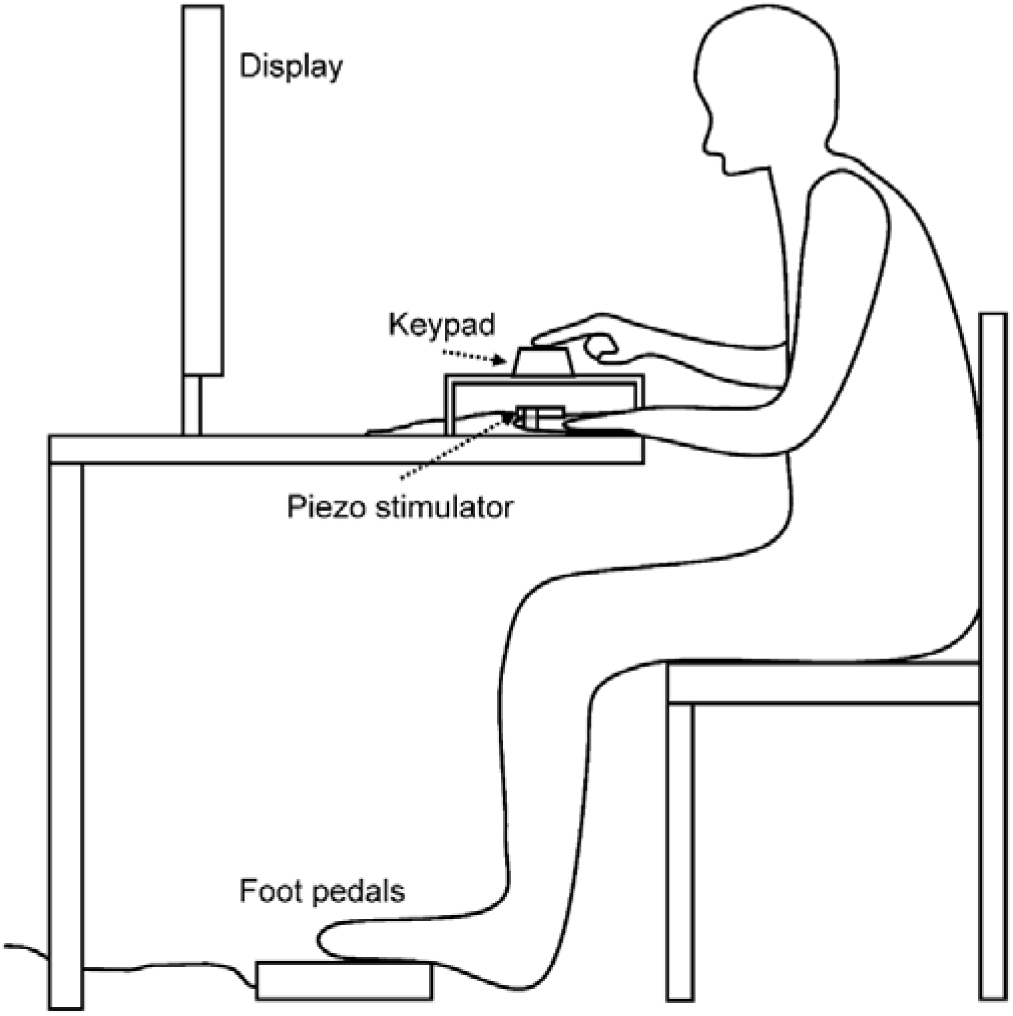
Experimental setup. Participants sat facing a display with their left hand placed on a table inside a box, palm facing up. A tactile stimulator was attached to their left index fingertip, and a keypad was positioned on the box, directly above the tip of the index finger. Participants were instructed to press the key with their right index finger to trigger the tactile stimulus during the task and to use the foot pedals to response for the intensity judgment.

The EEG was recorded with a BioSemi Active-Two amplifier system (BioSemi) from the AFz, F7, F3, Fz, F4, F8, FC3, FC1, FCz, FC2, FC4, C5, C3, C1, Cz, C2, C4, C6, CP3, CP1, CPz, CP2, CP4, P7, P3, Pz, P4, P8, PO7, PO3, POz, PO4, PO6, and Oz electrodes using Ag-AgCl active electrodes according to the international 10–20 system, mounted in a headcap. Six additional flat electrodes were attached to the left mastoid, right mastoid, outer canthi of both eyes, and above and below the right eye. Conducting gel was applied to the electrodes to ensure good contact with the skin. The offset potentials of all electrodes were kept below ±25 μV. EEG signals were recorded at a sampling frequency of 2048 Hz and referenced online against the CMS and DRL electrodes of the BioSemi system.

### Design and Procedure

Participants were tested individually in an electrically noise-shielded room. They were seated approximately 60 cm from the display. In each trial, participants were asked to press the button on the box with their right hand at a self-paced rhythm to trigger the tactile stimulus or to wait for the tactile stimulus (depending on the condition). In the no-delay condition of Experiment 1 and the active condition of Experiment 2, the 2-second target stimulus was delivered to the participant’s finger immediately after the key press. In the delayed condition of Experiment 1, the stimulus was delivered after a random delay of 500-1000 ms. In the passive condition, the stimulus was delivered 500-1000 ms after the start of the trial. The second tactile stimulus, referred to as the standard stimulus, was always delivered 500 ms after the target stimulus. Participants responded to the question of which of the two vibrations felt stronger using the foot pedals. During the stimulation, a white fixation dot was presented in the center of the display.

The intensity of the tactile stimulus was adjusted by modifying the height of the vibration pins. The height of the standard stimulus (i.e., the second tactile stimulus) was fixed at 1.12 mm, while the height of the target stimulus varied among participants. First, we measured the 50% discrimination threshold from the standard stimulus for each individual using the QUEST method (Watson, 2017; Watson & Pelli, 1983) in the pre-matching task. This threshold should theoretically feel equally strong as the standard stimulus when applied first. In the pre-matching task, we applied an upward staircase and a downward staircase, each with 30 trials. The initial amplitudes of the upward and downward staircases were 0.85 mm and 1.35 mm, respectively. The two tactile stimuli were 2 seconds long and were applied passively with an interval of 500 ms between them. Participants responded to the question of which stimulus felt stronger using the foot pedals. The amplitude of the first stimulus was updated in each trial according to the previous responses using the QUEST method. EEG recording was not conducted during the pre-matching task.

The individual 50% discrimination threshold acquired in the pre-matching task was used for the target stimulus in the main task. The trials using the target stimulus were repeated 30 times. To create variance in the main task, we added 18 dummy trials in which the amplitudes of the first stimulus were ± 0.05 mm, ± 0.1 mm, or ± 0.15 mm from the 50% discrimination threshold (each repeated 3 times). The dummy trials were excluded from the statistical and EEG analyses. The trial order was randomized for each participant.

In summary, participants first completed a pre-matching task to determine the individual intensity of the target stimulus. In the main task, they completed 30 trials containing the target stimulus and 18 dummy trials with slightly stronger or weaker stimuli. In both experiments, two 2-second tactile stimuli were always delivered sequentially, with a 500-ms interval. The initiation of the first stimulus varied depending on the experiment and condition. In Experiment 1, the no-delay and delayed trials were mixed and randomized. In Experiment 2, the active and passive trials were blocked because the actions participants needed to perform differed between these trials. The block order was counterbalanced among participants. Additionally, 6 practice trials were conducted before the pre-matching task in both experiments. Another 6 practice trials were conducted before the main task, containing one target trial, one +0.15 mm dummy trial, and one −0.15 mm dummy trial for each condition. Participants whose 50% discrimination threshold exceeded 1.5 mm (the maximum height of the pins) in at least one staircase were excluded from the main task. One participant was excluded from each experiment due to this criterion.

### EEG Data Processing

EEG signals were pre-processed using the MNE-Python package (Gramfort et al., 2013) in Python. A 1–200 Hz bandpass filter (basic linear finite impulse response filter) was applied to remove slow drifts and high-frequency noise from the EEG signals. The EEG signals were then re-referenced to the average of the left and right mastoids and segmented into epochs ranging from 0–2 seconds after the onset of the first vibration. Fast Fourier Transform (FFT) was conducted for the 2-second epoch, and signal-to-noise ratios (SNRs) were calculated. For the calculation of SNR, the maximum amplitude within the 71±6 Hz window was divided by the average amplitude in this window, excluding the peak bin.

To further analyze the temporal dynamics of the SSSEP, we applied a sliding window analysis. A 1-second window was used. Because FFT results in a spectral resolution inversely proportional to the window length, the 1-second length is the minimum required to calculate the SNR for the target frequency. The time window was slid every 100 ms, resulting in 11 time windows. Additionally, to confirm that the SNR did not differ between conditions before the target stimulus, another time window from −1 to 0 seconds before the onset of the target stimulus was analyzed.

## Results

### Experiment 1: No-delay vs delayed conditions

Figure 2 shows the ratio of selecting the first tactile stimulation as stronger than the second one. The larger this ratio, the stronger participants perceived the first (target) touch to be. The selection ratio for the no-delay condition was significantly larger than that for the delayed condition (*t*(17) = 3.08, *p* = .007, d = 0.73). Thus, participants perceived the no-delay touches as stronger than the delayed touches, which is the opposite of the typical sensory attenuation phenomenon.

**Figure 2.**
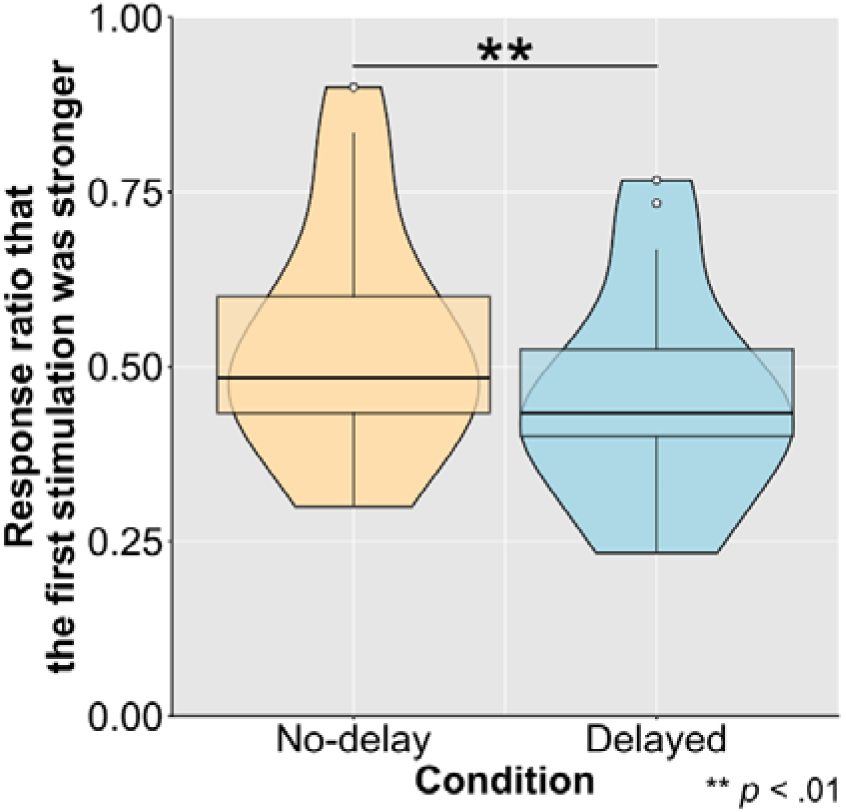
Response ratio for selecting the first stimulation as the stronger one in Experiment 1. The boxes represent the interquartile range (IQR, the middle 50% of the data). The horizontal line inside the box marks the mean. The whiskers represent the smallest and largest value within 1.5 times the IQR. The violin plot visualizes the central tendency and dispersion of the data. Data points that fall outside 1.5 times the IQR are plotted as white circles. Asterisks denote significant differences between the two conditions (*p* < .01). The same applies for other box-violin plots in Figure 3 and 5.

Next, Figure 3A shows the SNRs of the entire 2-second target stimulus for the no-delay and delayed conditions at C2, and Figure 3B shows the topography of the difference between the no-delay and delayed conditions. Across all electrodes, the SNRs were significantly greater in the no-delay condition than in the delayed condition (Bonferroni method was applied for multiple comparisons; *t*s (17) > 2.82, *p*s < .02, ds > 0.66). These results indicate that the neural response was overall larger for no-delay touches than for delayed touches.

**Figure 3.**
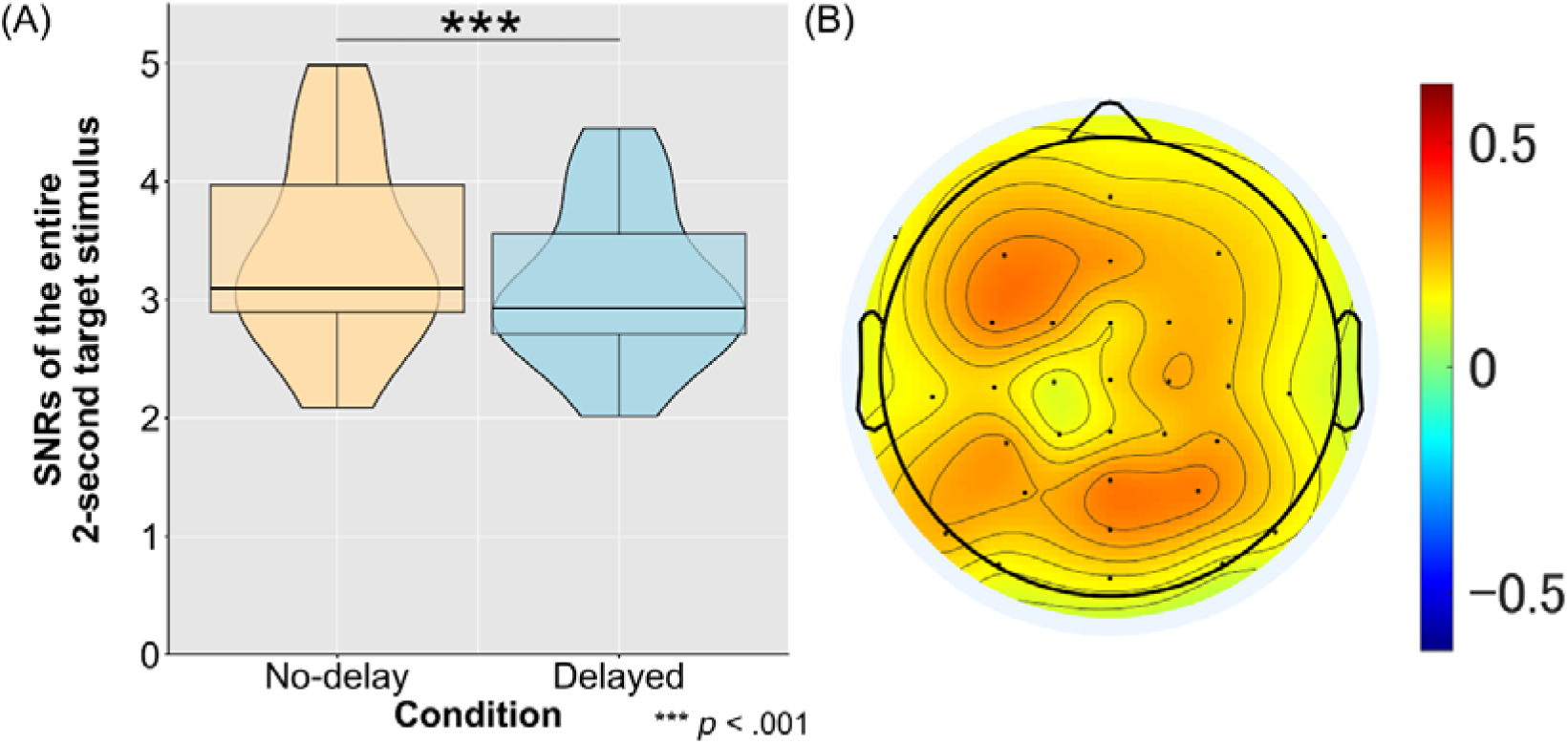
SNRs and topography for the entire 2-second tactile stimulus in Experiment 1. The SNR of the SSSEP significantly differed between the no-delay and delayed conditions (A). The topographical map of the SNR difference between the no-delay and delayed conditions is shown in (B). Higher values (red) indicate that SNRs in the no-delay condition are greater than those in the delayed condition.

Furthermore, Figure 4A shows the SNRs in the 11 one-second time windows, each slid by 100 ms, at C2. The SNR of another one-second time window before the onset of the target stimulus (i.e., −1 to 0 s) was also calculated and added to the figure to confirm that the baseline between the no-delay and delayed conditions did not differ. Figure 4B shows the topography of the *t* values when comparing between the no-delay and delayed conditions across the 12 time windows. We conducted *t*-tests to compare the difference between the no-delay and delayed conditions at C2. The significance level was set to .004 (i.e., 0.05/12) according to the Bonferroni correction. There was no significant difference between the two conditions before the onset of the stimulation (*t*(17) = 0.26, *p* = .80, d = 0.06). For the time window of 0-1000 ms, the amplitude of SSSEP in the no-delay condition was significantly smaller than in the delayed condition (*t*(17) = 4.23, *p* < .001, d = 1.00), consistent with the phenomenon of sensory attenuation, indicating a smaller neural response for the no-delay touch than for the delayed touch. However, this phenomenon disappeared in the second and third time window (*ts*(17) < 3.07, *p* > .08, d =.72) and reversed from the time window of 300-1300 ms and in all following windows (*t*s(17) > 4.25, *p*s < .001, ds > 1.00). Similar results were observed across almost all electrodes (Figure 4B).

**Figure 4.**
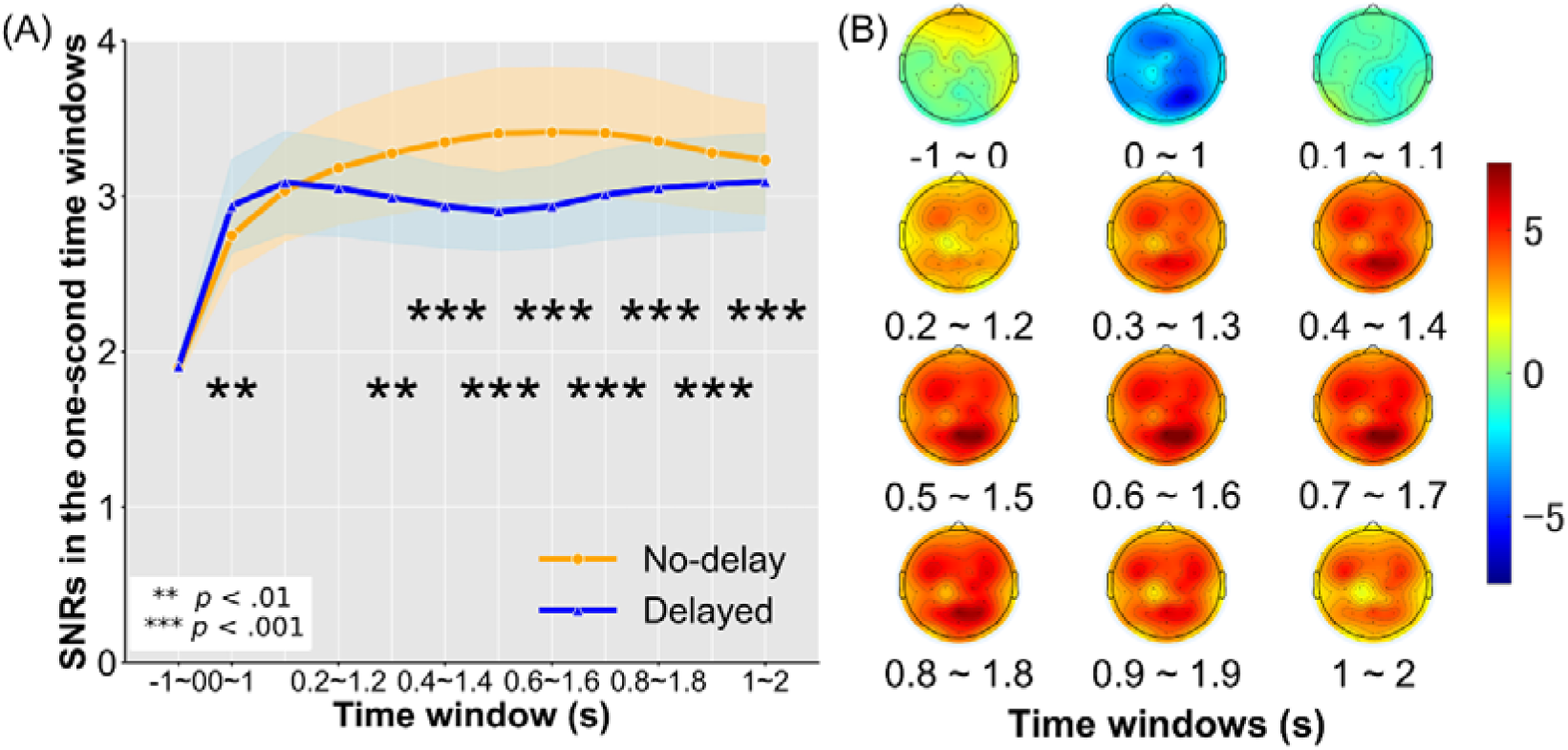
SNRs at C2 and topographies of *t* values in each time window in Experiment 1. (A) shows the temporal changes in SNR for each condition across the 11 time windows, as well as an additional window before the onset of the stimulus (i.e., −1 to 0 s) at C2. Shaded areas represent the 95% confidence interval. Asterisks indicate significant differences between the no-delay and delayed conditions. (B) shows the topographical maps of *t* values for the SNR difference between the no-delay and delayed conditions in each time window. Positive values (red) indicate that SNRs in the no-delay condition are greater than in the delayed condition, while negative values (blue) indicate the opposite.

### Experiment 2: Active vs passive conditions

Experiment 2 compared the subjective judgment of intensity and SSSEP between the active and passive conditions. The response ratio did not significantly differ between the active and passive conditions (*t*(15) = 0.89, *p* = .39, d = 0.22; Figure 5). On the other hand, the pattern of SSSEP amplitude replicated the findings from Experiment 1. First, for the overall SSSEP amplitude of the entire 2-second trial, the SSSEP amplitudes were significantly greater in the active condition than in the passive condition across all electrodes (Bonferroni method was applied for multiple comparisons; *t*s(15) > 2.39, *p*s < .03, ds > 0.60). Next, for the SSSEP in each time window at C2 (Figure 6A), the SNR did not differ before the onset of the stimulation between the active and passive conditions (−1 to 0s; *t*(15) = .95, *p* = .36, d =.24). For the time window of 0-1000 ms, the SNR in the active condition was significantly smaller than in the passive condition (*t*(15) = 4.36, *p* < .001, d = 1.09), reflecting an attenuated neural response for self-generated sensory input compared to externally generated input. This attenuation disappeared in the next time window (100-1100 ms; *t*(15) = .91, *p* = .38, d = .23) and reversed from the time window of 200-1200 ms and in all following windows (*t*(15) > 3.36, *p* < .05, d > .84). This temporal dynamic was consistent across all electrodes (Figure 6B).

**Figure 5.**
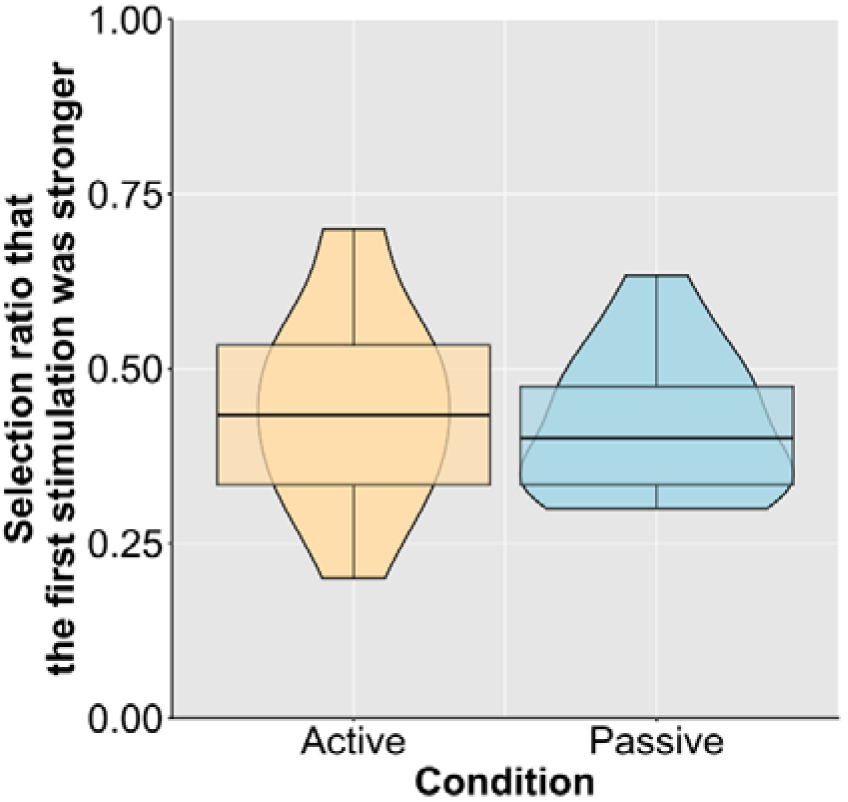
Response ratio for selecting the first stimulation as the stronger one in Experiment 2. There was no significant difference in the subjective intensity judgment between the active and passive conditions.

**Figure 6.**
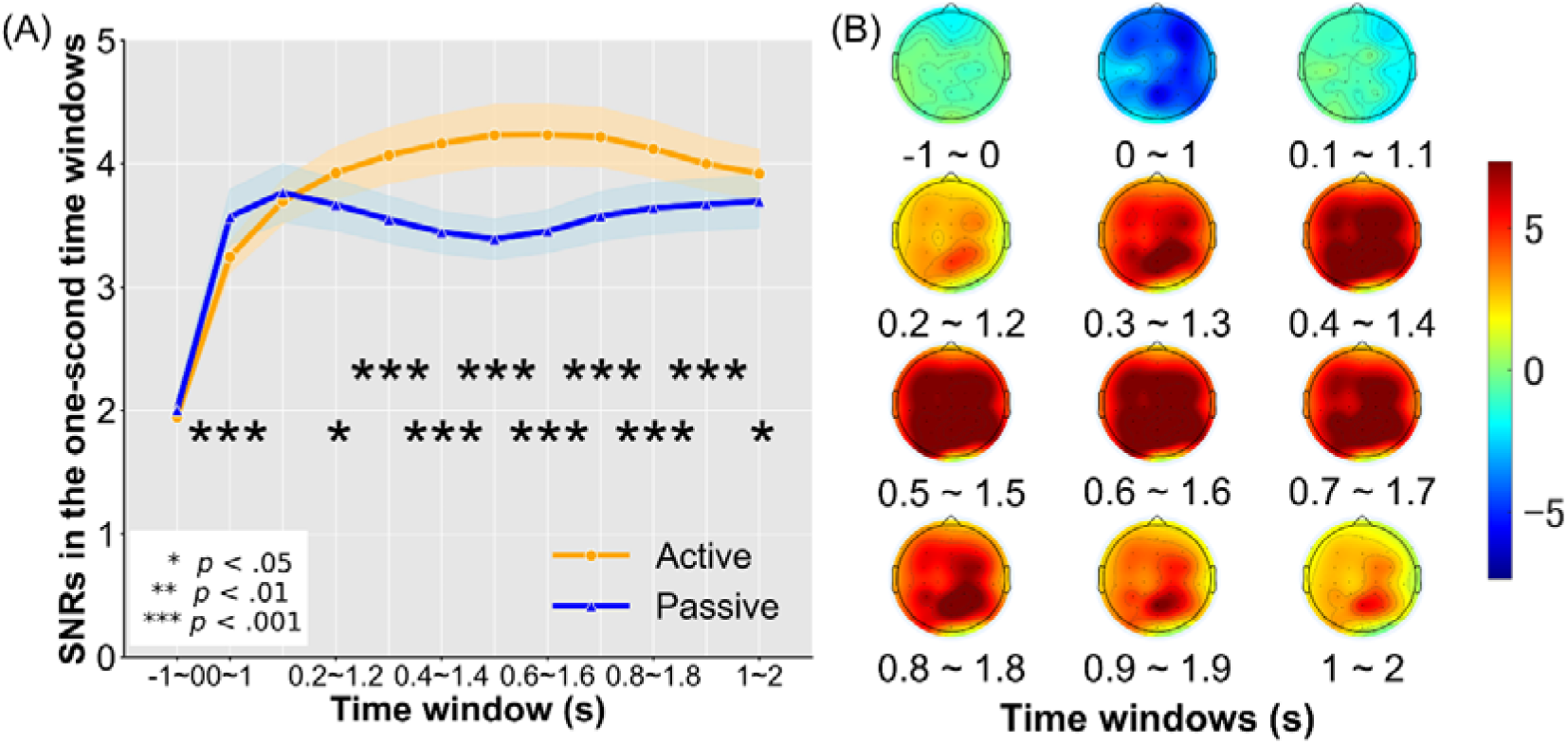
SNRs at C2 and topographies of *t* values in each time window in Experiment 2. (A) shows the temporal changes in SNR for each condition across the 12 time windows at C2, and (B) shows the topographical maps of *t* values for the SNR difference between the active and passive conditions in each time window. Positive values (red) indicate that SNRs in the active condition are greater than in the passive condition, while negative values (blue) indicate the opposite. The temporal dynamics of the SSSEP replicated the findings from Experiment 1, showing an early sensory attenuation followed by a later sensory enhancement for voluntary actions.

## Discussion

Sensory attenuation is a well-known mechanism for distinguishing between self-generated and external sensory input. Specifically, when our actions cause sensory inputs, the sensory predictions from efference copies of our motor commands result in a cancellation of sensory processing. This mechanism allows us to recognize which sensory input is caused by ourselves and which is externally generated. However, behavioral reports have shown both attenuation and enhancement for self-generated sensory input (van der Weiden et al., 2015; Yon et al., 2021; Yon & Press, 2017). Self-generated sensory input can be predicted and may therefore sharpen our perception (Press et al., 2020; Yon et al., 2018). This paradox remains largely unresolved. The present study used SSSEP and sliding window analysis to examine the temporal dynamics of neural responses to self-generated tactile stimuli compared to delayed stimuli (Experiment 1) or externally generated stimuli (Experiment 2). Our results showed that the self-generated stimuli were perceived significantly stronger than the delayed stimuli behaviorally (Experiment 1). The SSSEP results demonstrated sensory attenuation in the first time window (i.e., 0-1 s) for self-generated tactile stimuli compared to delayed or externally generated stimuli. However, this effect disappeared quickly and was later replaced by a significant enhancement effect for self-generated tactile stimuli. These findings suggest that sensory input caused by our own actions is rapidly attenuated in the very early phase but is enhanced in the later phase.

Sensory attenuation is often introduced by citing the well-known phenomenon that we cannot tickle ourselves because our brain predicts sensory input caused by our own actions (Blakemore et al., 2000). The attenuation theory also aligns with the observation that patients with schizophrenia can tickle themselves and often have difficulty distinguishing between self-generated and externally generated sensations (Blakemore et al., 2000; Frith et al., 2000; Lemaitre et al., 2016). However, sensory predictions can potentially link to preactivations in our perceptual networks (Roussel et al., 2013), enhancing perceptual processing for sensory input that matches the predictions (Wen et al., 2018). Furthermore, Bayesian theories suggest that anticipations should enhance perceptual processes rather than suppress them (Press et al., 2020). Our findings suggest that this enhancement from anticipated sensory input does indeed occur, but it happens later in the process, following an initial phase of attenuation. The early attenuation is likely automatic and crucial for recognizing self-generated sensory input without the need for explicit judgment. With such online sensory gating mechanism during movements, the brain can avoid overwhelming itself with self-caused sensory noises (Angel & Malenka, 1982; Chapman et al., 1987; Collins et al., 1998; Milne et al., 1988; Wolpert & Flanagan, 2001). Since humans constantly interact with the world – walking, taking, thinking – each movement generates sensory input. The implicit and automatic sensory attenuation mechanism allows the brain to suppress unnecessary sensory processing for constant actions. For such purpose, this mechanism requires fast and accurate temporal dynamics.

In contrast, the later enhanced neural activity, observed in the later time windows and onward in our SSSEP recording, is likely linked to higher-level and more informative processing. When we interact with the environment with goal-directed actions, self-generated sensory input is often meaningful, and our perceptual network likely allocates more resources to this input to enable more accurate perceptual processing and detections of self-caused events. There are increasing evidence showing that voluntary actions benefit perceptions. For example, a previous study reported that voluntary actions lowered the hearing threshold for self-generated sounds, and brain activity in the auditory cortex was enhanced in the hemisphere contralateral to the active hand (Reznik et al., 2014). Another study reported that when the motion of a visual stimulus was consistent with participants’ own hand movements, the dominance duration of that visual stimulus was extended in a visual competition paradigm (Maruya et al., 2007). In summary, the detection of self-generated sensory input is likely to benefit from the predictions based on motor commands. Furthermore, it is suggested that attentional mechanisms may play a role in modulating brain responses to self-generated and externally generated sensory input (Ackerley et al., 2012). It is known that people use expectations to guide temporal attention (Duyar et al., 2024). The temporal dynamics of attention allocation are considered to increase until 500 ms (Moon et al., 2019). This is much slower and longer than the sensory attenuation mechanism. In summary, the sensory attenuation and sensory enhancement to self-caused events may serve different functions in humans and likely have distinct temporal dynamics. This also align with hierarchical predictive coding, which suggests that expectations play different roles in early and later sensory processing (Chennu et al., 2013).

Lastly, sensory attenuation is often reported in behavior results. However, we only observed sensory enhancement behaviorally in the present study, likely due to the characteristics and length of our tactile stimulation. The subjective report of intensity was consistent with the overall neural response to the stimulation. Steady-state evoked potential (SSEP) measures require repetitive high-frequency stimulation, which may differ significantly from the sensory input we typically experience. Nevertheless, SSEP measures allow us to examine neural responses during continuous sensorimotor processing (Norcia et al., 2015; Severens et al., 2010), providing valuable insights into the temporal dynamics of sensorimotor processing when combined with sliding time window analysis. Additionally, SSEP is advantageous for studying populations with difficulties in subjective reporting, such as infants and children, and may be useful for investigating impairments in sensory predictions in psychosis (Bansal et al., 2018; Roussel et al., 2013).

In summary, the present study utilized SSSEP to investigate the temporal dynamics of perceptual processing for self-generated sensory input. We identified an early sensory attenuation followed by a later sensory enhancement. Humans use sensory attenuation to distinguish between self-generated and external inputs and suppress sensory noises from their own actions in the early phase, while also allocating more cognitive resources to self-generated input later on to enhance perceptual processing.

## Acknowledgement

The authors thank Souta Hidaka for the valuable discussion.

## Funding

This work was supported by JST FOREST Program (Grant Number: JPMJFR2144) and JST Moonshot R&D Program (Grant Number: JPMJMS2013).

## Author Contributions

YS: conceptualization, data collection, analysis, writing – original draft; AC: conceptualization, analysis, writing – original draft; SH: conceptualization, writing – review & editing; WW: conceptualization, analysis, writing - original draft.

## Notes

### Competing Interest Statement

The authors have declared no competing interest.

